# Recapture rates and habitat associations of White-faced Darter *Leucorhinnia dubia* on Fenn’s and Whixall Moss, Shropshire, UK

**DOI:** 10.1101/349936

**Authors:** Rachel Davies, Achaz von Hardenberg, Matthew Geary

## Abstract

Land-use change and habitat loss are important drivers of biodiversity decline at both global and local scales. To protect species from the impacts of land-use change it is important to understand the population dynamics and habitat associations across these scales. Here we present an investigation into the survival and habitat preferences of White-faced Darter (*Leucorrhinia dubia*) at the local scale at Fenn’s and Whixall Moss, Shropshire, UK. We used capture-mark-recapture methods to investigate survival and used sightings of individual dragonflies along with habitat data to investigate habitat preference. We found that survival between capture-visits was very low and that White-faced Darter show a clear preference for the open moss habitat on this site. In both cases, we found that the detectability either through sightings or recaptures was potentially very low and suggest that this should be taken into account in future analyses. We suggest that this could be achieved by encouraging recorders to submit complete lists and to repeat visits to sites.

## Introduction

There has been a marked decline in global biodiversity in the last several decades, a decline which is expected to continue, and this has been largely attributed to changes in land-use activities (Sala, 2000). Land-use activities include agriculture, forestry, creation of urban areas, and use of natural resources (Foley et al., 2005). These activities have a huge impact on environmental characteristics and often cause habitat loss and fragmentation, contributing largely to the decline in global species diversity (Holloway et al., 2003). As such management and protection of habitats and populations is vital at both a local and global scale (Foley et al., 2005; Holloway et al., 2003).

A bias exists in conservation research towards charismatic vertebrates (Di Marco et al., 2017). Although Odonata are charismatic invertebrates they are not immune to this bias (Clausnitzer et al., 2009). In addition much research into Odonata focuses on physiology, evolution and behaviour (Córdoba-Aguilar, 2008) and they have rarely been the focus of conservation research (Clausnitzer et al., 2009). Basic ecological research into demography, survival and habitat use is essential for effective protection of species and habitats. For any taxa this requires detailed ecological and life history data collected in the field. These are often difficult to obtain, particularly on large scales. Integrating large scale data such as presence-only distribution datasets with more detailed local information is a current challenge in conservation ecology (Powney and Isaac, 2015).

Methods to analyse habitat preferences are varied depending on the data available. The current ‘gold standard’ is the use of site occupancy models which take into account detectability (i.e. the probability that a species is detected in a site if present) when estimating occupancy: the probability that a species is present in a site (MacKenzie et al., 2003). Models using this framework help us to avoid the age-old problem of “imperfect detection”, i.e. failing to spot a species during a survey on a site where it is actually present (MacKenzie et al., 2003). However, these models require repeated surveys where both detections and non-detections are recorded; these data are not always available, especially in the case of historic data collected by volunteers. On larger scales a number of methods exist which can use only presence records along with environmental covariates (Elith and Leathwick, 2009). These can tell us about habitat use but are constrained to estimate a measure of the relative importance of habitats rather than the true probability of presence (Elith et al., 2011) and are limited by the environmental data available. At very small scales, such as individual protected areas, detailed data on habitats and landcover can be difficult to obtain. Datasets such as the UK landcover map (LCM2015), although the resolution is 15m, are too crude for local studies in some areas. Simpler methods which indicate preferred habitat, such as selection indices (Manly et al., 2007), have fewer assumptions and can be revealing even at small scales (Neu et al., 1974).

Investigating survival and movement requires individual recognition and methods using a capture-mark-recapture approach are well established (McCrea and Morgan, 2014). Such analyses can tell us about the age-sex specific survival probabilities of individuals, the use of different sites or habitats and how these change over time, and the likelihood of encountering individuals again in the future. High quality data of this type can provide accurate estimates of population size. Capture-mark-recapture methods have been used on Odonata populations in the past to monitor rare species (Cordero-Rivera and Stoks, 2008) as well as to study the life history of more abundant species (Anholt et al., 2001). Odonata, because of the ease by which they can usually be individually marked, have also been used as model species for methodological research on the development of capture-mark-recapture techniques (Manly and Parr, 1968).

The White-faced Darter (*Leucorrhinia dubia*), is a specialist of lowland peatbogs where it breeds in bog pools containing sphagnum mosses (Smallshire and Swash, 2014). It has a life cycle that includes a 1-3-year larval period, followed by an adult flight period (Smallshire and Swash, 2014). Emergence is weather dependent and will typically start in either May or June each year. Tenerals are thought to disperse to low scrub following emergence, staying there whilst they mature. Following this, the adults return to breeding pools, with males returning sooner than females so they can hold breeding territories (Smallshire and Swash, 2014). The adult flight period typically ends in either late July or August. The White-faced Darter has a scattered distribution and its populations have been declining in Britain over the past several decades. Despite being classified as a species of least concern on the IUCN Red Data List (Clausnitzer et al., 2009), this decline in Britain has resulted in a classification of Endangered on the Odonata Red Data List for Britain (Daguet et al., 2008). This decline is largely attributed to habitat loss and the resulting habitat fragmentation (Daguet et al., 2008); as over 90% of England’s peatbogs have been lost or substantially damaged to date (English Nature, 2002). There are currently only three stable historical populations of White-faced Darter in England, along with two recently reintroduced populations, one in Cumbria and one in Cheshire (Clarke, 2014; Meredith, 2017).

Here we use two methods to investigate important ecological characteristics of White-faced Darter on Fenn’s and Whixall moss in Shropshire, UK. We use capture-mark-recapture methods to investigate survival and movements of adults during the flight period and a selection index method to investigate habitat use. These methods can both contribute to our understanding of the spatial use of habitat by White-faced Darter and can help us to prioritise future research for this species.

## Methods

### Study area

Fenn’s, Whixall and Bettisfield Mosses (FWB Mosses) are located within Shropshire (52?55ϏN 2°46ϏW) and they support a large, long-established population of White-faced Darter. FWB Mosses are a lowland raised bog complex, stretching nearly 1000 hectares (Meredith, 2017). Historically, the mosses were used for peat cutting and in the 19^th^ century they were drained to allow larger-scale operations to take place (Meredith, 2017). Eventually, in 1990, the mosses were taken over by English Nature (now Natural England) and long-term restoration began, benefitting a whole host of mossland species, including the White-faced Darter (Meredith, 2017).

### Field methods

The site was surveyed twice per week between 22 May and 6 July 2017. This encompassed the peak flight period of White-faced Darter (Smallshire and Swash, 2014). Two separate breeding pools within FWB Mosses were sampled simultaneously, along with a variety of scrub and other potentially suitable habitat. On each sampling occasion, the full sampling area was searched for any White-faced Darter individuals. Different routes were walked on each occasion to allow different areas within the sampling area to be searched at different times of the day. Sampling sessions lasting between 5-10 hours, being carried out between 10am and 4pm, as this is the favoured flight period for adult dragonflies (Smallshire and Beynon, 2010). Sampling days were weather dependent (Chin and Taylor, 2009) and weather conditions were recorded on all sampling days.

#### Capture-Mark-Recapture

Where possible, mature adults were caught using an invertebrate net and marked with a unique number on their wing (Chin and Taylor, 2009), using an Edding 404 permanent marker pen (Plate 1). The insects were then released at point of capture and any behavioural observations recorded. Not all observed individuals were captured and tenerals were excluded from the capture-mark-recapture survey as during this life stage they are fragile and handling may cause wing damage (Allen and Thompson, 2010). Tenerals are easily identified by their pale green colouration, a lack of their full adult colouration and by their shiny wings (Smallshire and Swash, 2014). Insects recaptured on day of marking were not re-counted (Foster and Soluk, 2004). Following an initial marking, recapture on successive days was only necessary when relevant information could not be collected from re-sighted individuals (Lettink and Armstrong, 2003).

**Plate 1:**
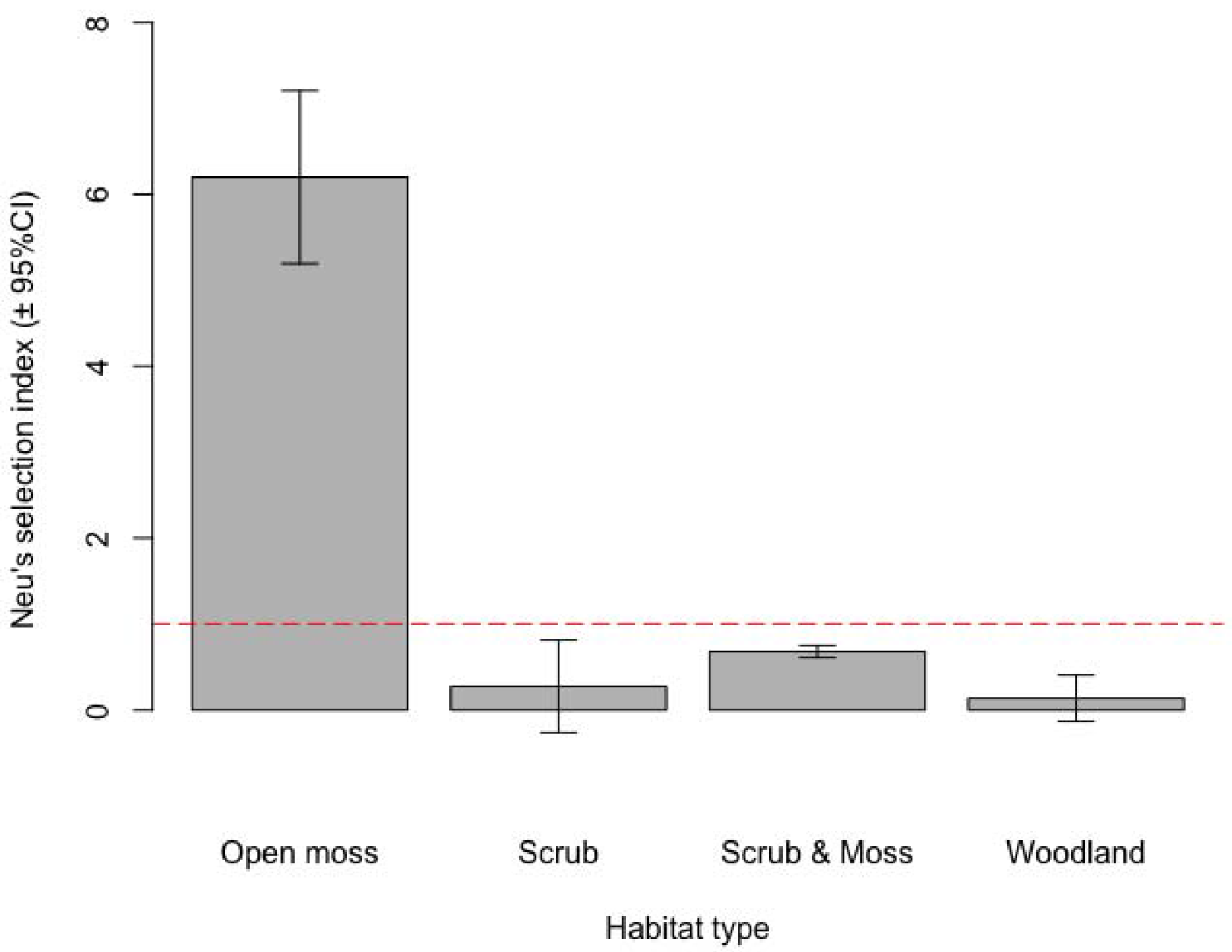
A marked male White-faced Darter at Fenn’s and Whixall Moss in 2017.

#### Habitat selection index

White-faced Darter presence was recorded while searching the site during the capture-mark-recapture study. This included captured individuals as well as those seen on survey routes but not captured. On each occasion the location of the individual was recorded with a hand-held GPS unit (Garmin GPSMAP 64). Additionally, a phase 1 habitat survey (Joint Nature Conservation Committee, 2010) was conducted across the study site to produce a habitat map using 100 × 100 m grid cells. The proportions of five habitat types were recorded in each square: moss (peat moss, rushes and sedges), scrub (low woody vegetation), scrub-moss (peat moss with low woody vegetation), water (open pools) and woodland (mature trees). From this the dominant habitat in each square was calculated. Of these, only water was not used in analyses as adult individuals tended to be sighted over terrestrial habitat.

### Data analysis

#### Capture-mark-recapture

We estimated the daily survival probability and the probability of recapture using a continuous-time open capture-mark-recapture model as described in Fouchet et al., (2016). This model relaxes the discrete-time assumption of classic capture-mark-recapture models, allowing robust estimates also in the case of lags between capture sessions of varying duration. The analysis was carried out using the *CMRT* package (Santin-Janin and Fouchet, 2015) in R version 3.5.0 (R Core Team, 2018).

#### Habitat selection index

Selection indices calculate habitat use as a ratio between habitat where a species is recorded compared to the proportion of each habitat within the study area (Manly et al., 2007). Although relatively simple they can be effective in indicating habitat use (Manly et al., 2007). Selection indices can be sensitive to the scale used in calculating habitat use however Neu’s index is relatively robust to changes in scale, (Neu et al., 1974). For this reason we used Neu’s index which calculates 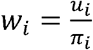 where *w*_*i*_ is the proportion of squares of each dominant habitat type among all of the squares with White-faced Darter records and *π*_*i*_ is the proportion of each dominant habitat type among all of the squares in the study area. Values of the index 1 indicate use of a habitat type in greater proportions than it is generally available in the study area. Selection index analysis was performed in R version 3.5.0 (R Core Team, 2018) using the *adehabitatHS* package (Calenge, 2006).

## Results

### Capture-Mark-Recapture Model

A total of 13 sampling days were carried out at FWB Mosses from the 22^nd^ May 2017 until the 7^th^ July 2017. During these sampling days, a total of 50 adult White-faced Darters were marked (41 males and 9 females), and a total of 6 recaptures were made. Probability of survival between sampling days was estimated at 0.06 (95% confidence intervals: 0.02-0.17). Probability of recapture on each sampling day was estimated at 0.05 (95% confidence intervals: 0.00-0.11). Further models using a range of co-variates were unsuitable as the models became over parameterised due to the lack of recaptures.

### Selection Index

A further 304 individual White-faced Darter were observed during the fieldwork, 234 of which were not captured but only observed from distance (Figure 1). White-faced Darter show a clear preference (Neu’s index > 1) for ‘moss’ habitats while ‘scrub’, ‘scrub and moss’ and ‘woodland’ (Neu’s index < 1) appear to be avoided (Figure 2).

**Figure 1:**
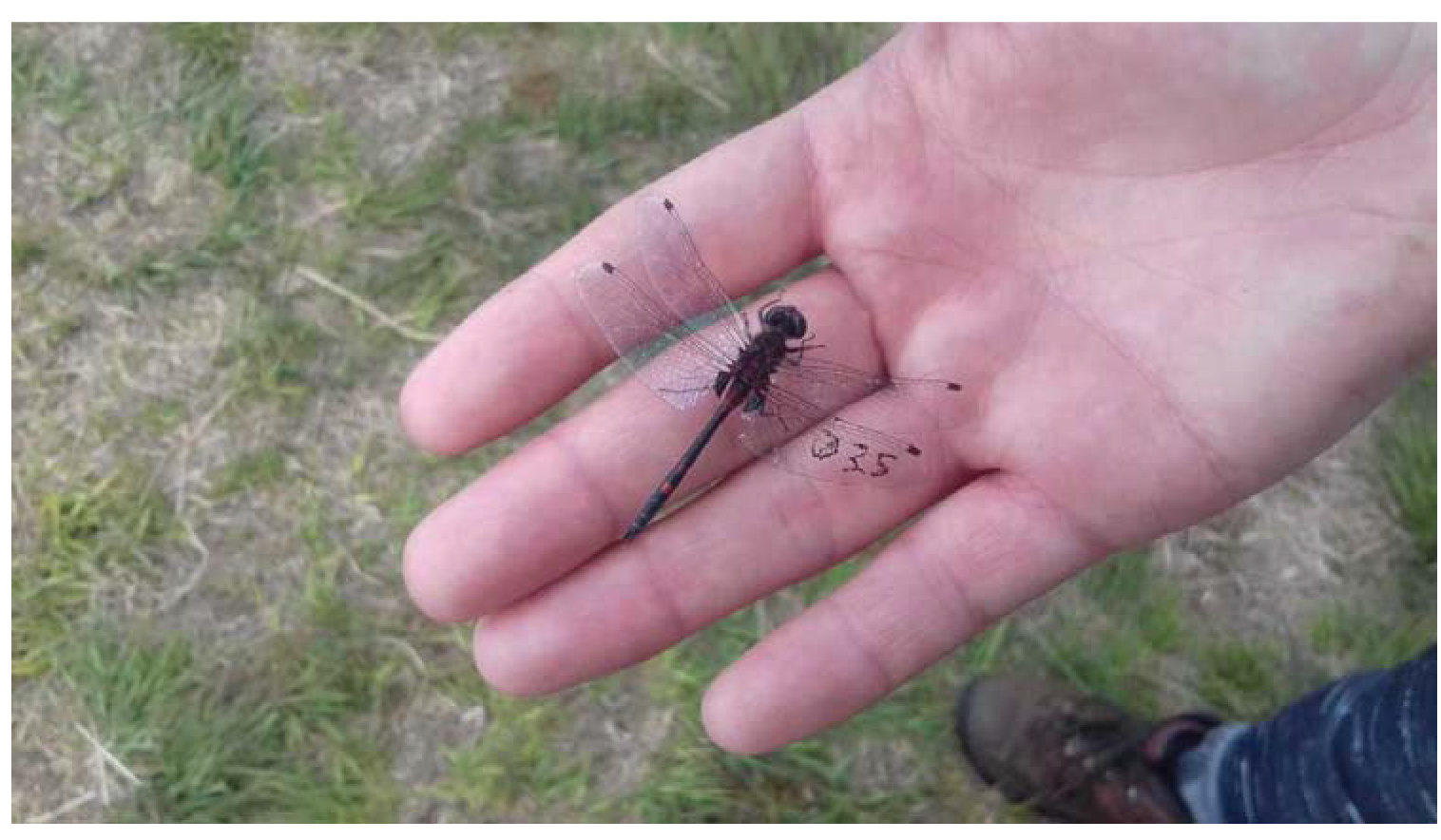
Number of White-faced Darters of different sex and age classes recorded in Fenn’s and Whixall Moss in May-July 2017.

**Figure 2:**
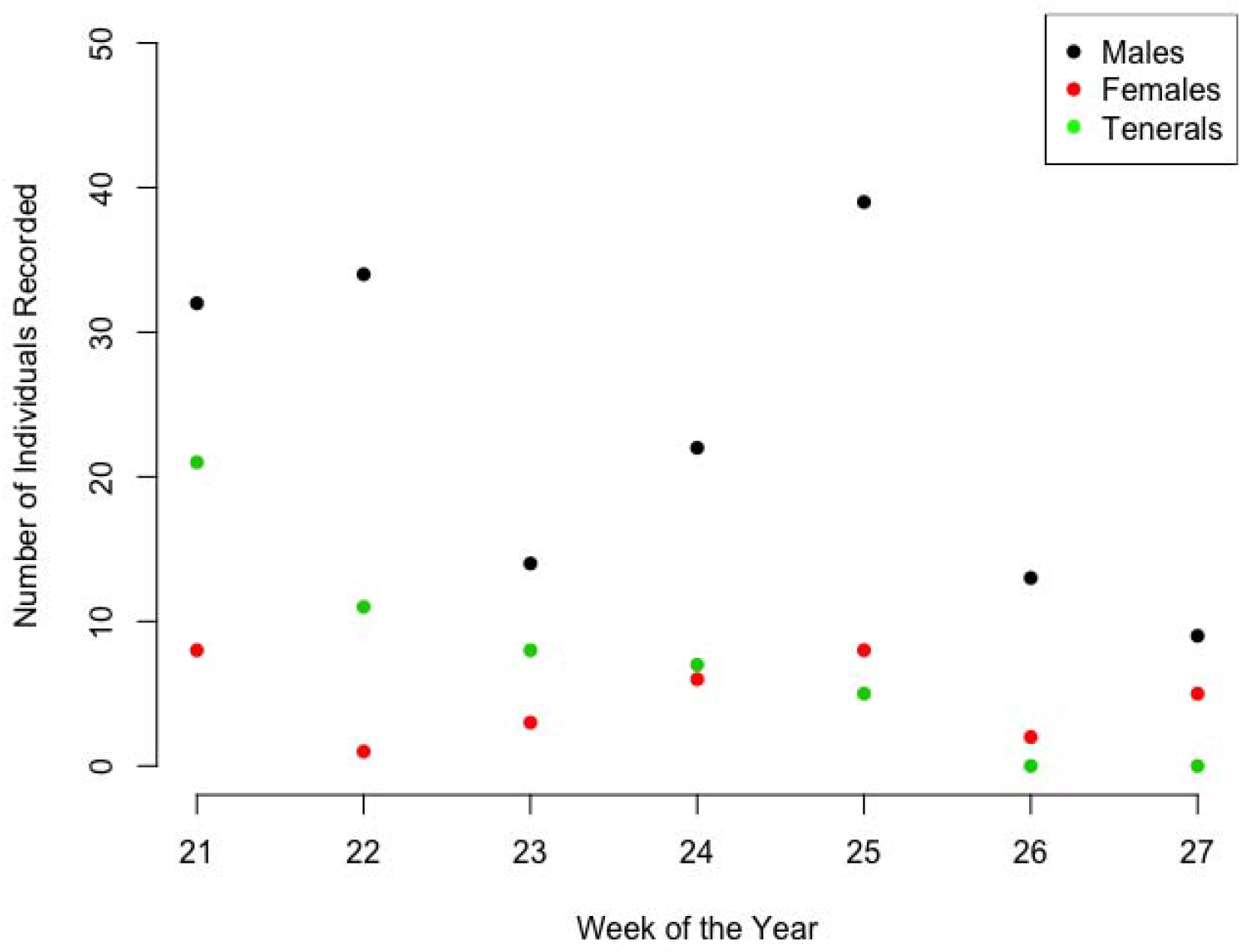
Neu’s selection index for White-faced Darter records on Fenn’s and Whixall Moss. The red dashed line represents a selection index of 1 (i.e. no selection). If the confidence intervals do not include the red line and are above it, the habitat is positively selected, while if the confidence intervals stay below the red line, the habitat is negatively selected.

## Discussion

The capture-mark-recapture model suggested that the survival rate of adult White-faced Darter from one survey session to the next was only between 2% and 17% (95% confidence intervals). Similarly, we had a very low recapture rate (only 6 recaptures in 13 days) and our capture-mark-recapture model estimated a 0-11% chance of each individual being recaptured in successive capture sessions. Although low capture rates might be expected in a large invertebrate population and have been noted before in Odonata (Cordero-Rivera and Stoks, 2008), this was lower than expected. Although male White-faced Darter hold territories they are less tied to these sites than species such as Four-spotted Chaser (*Libellula quadrimaculata*) and so are less predictable in their movements (Merritt et al., 1996). We suggest that future capture-mark-recapture approaches for this species, and other similarly cryptic species, need a greater number of capture days and more researchers in the field making captures. This increase in effort is likely to increase the capture rate and increase the accuracy of estimates.

Many more White-faced Darter were seen than were captured and the resource selection index calculated using these data suggest that they prefer the ‘moss’ habitat among those available. Although this habitat is the most common habitat in the study area, the selection index suggests that they use this habitat in proportions greater than those available across the site. The ‘moss’ habitat consists of peat with low heather vegetation and wet flushes and is the habitat most commonly found at pool edges. This is the habitat described in previous research on White-faced Darter (Dolný et al., 2018) and described in Boudot and Kalkman (2015) including “peat moss, rushes and sedges”. Locally on this site, White-faced Darters appear to avoid complex vegetation, including scrub and woodland. However, White-faced Darter sites, especially those in Scotland which represents the stronghold for this species in Britain, are often forested (Cham et al., 2014). Breeding pools within these sites are likely to be in open areas but the association with woodland, particularly ancient woodland (Cham et al., 2014), is suggestive of some associations between White-faced Darter and these habitats at larger scales.

Moss habitats are certainly suitable for White-faced Darter, however, the low capture and recapture rate we found in this study may explain why open ‘moss’ habitats appeared to be preferred by this species. White-faced Darter are well camouflaged within their habitats and, as such, there is a good chance of missing individuals because of habitat complexity (i.e. low detectability, Mazerolle et al., 2007). Unfortunately, our field methods did not allow us to estimate detectability in terms of sightings on this occasion but the low capture probability found in our capture-mark-recapture study suggests it is very low. In future we suggest that survey methods are designed so that detectability can be estimated explicitly, in order to get more accurate estimates of occupancy and thus of resource selection. In this case we are left unable to confidently suggest whether White-faced Darter are avoiding more complex vegetation or whether they are harder to see and therefore record in these habitats.

Data which allows the estimation of detectability can easily be collected with just a few minor changes to currently common survey methods. In fact, the majority of these suggestions are already being requested by the British Dragonfly Society to provide data for the upcoming State of the Nation’s Dragonflies in 2020. We would like to add our voice to these calls to record complete lists and to repeat site visits. Complete lists are records of all the Odonata species detected on a single visit and allow non-detection to be inferred where species are not recorded (Isaac and Pocock, 2015). This requires recorders to note very common species as well as rarities. Unfortunately, there is a tendency in biological recordings to note only the rare or exciting species (e.g. first record of the year) and this can bias our inferences about population change amongst more common species (Isaac et al., 2014). Repeated site visits allow us to estimate the detectability of a species (MacKenzie et al., 2003) and consequently get unbiased estimates of occupancy, not affected by imperfect detection. We would also like to suggest that where possible recorders include some measure of effort in their surveys (e.g. time spent surveying or distance walked). Ideally this would be standardized and included in official protocols such as those already commonly in use for bird surveys.

We present the results in this paper as an indication of what can be done in terms of conservation research in Odonata. Although we have been unable to make firm inferences regarding White-faced Darter survival and habitat preference at this stage, due to the limited data, this study can provide valuable information which can contribute to the design of future studies. We suggest that research into the conservation ecology of White-faced Darter along with other Odonata species threatened with declining ranges, declining populations or habitat loss is essential to the long-term conservation of these species. Methods for such studies can be well informed by current practices used with other taxa. In particular, the analytical advances made in ornithology, research on Lepidoptera and work related to the use of data collected through citizen science provide a fantastic opportunity to advance our knowledge on the conservation ecology of Odonata.

## Acknowledgements

We would like to express our thanks to the British Dragonfly Society who supported this work financially, Natural England who provided permission to access the field site and to Chris Meredith who provided invaluable help in the field and advice on White-faced Darter ecology and behaviour. We would also like to thank all of the volunteers who helped with fieldwork in the summer of 2017.

